# Bacteriophage PAK_P3 genome structuration and dynamics during infection of *Pseudomonas aeruginosa* reveal specific interactions patterns

**DOI:** 10.64898/2025.12.15.694284

**Authors:** Amaury Bignaud, Quentin Lamy-Besnier, Devon E. Conti, Agnès Thierry, Fabien Girard, Pauline Misson, Romain Koszul, Laurent Debarbieux, Martial Marbouty

**Affiliations:** Institut Pasteur, Université Paris Cité, Spatial Regulation of Genomes Group, CNRS UMR 3525, Paris F-75015 France; Sorbonne Université, Collège Doctoral, Paris, France; Institut Pasteur, Université Paris Cité, CNRS UMR6047, Bacteriophage Bacterium Host, F-75015 Paris, France; Department of Biochemistry, University of Oxford, United Kingdom

## Abstract

Bacteriophages, or phages, are highly abundant and diverse genomic entities that play an important role in microbial ecology, evolution, and horizontal gene transfer. While the dynamic changes in genome organization in cellular organisms have been well described, the 3D folding of phage genomes during the infection of their host is extremely limited. Understanding how phage genomes fold, invade and rearrange the genome of their host to functionally organize the optimal expression of their genes remains unknown. Here, we explore the spatial dynamics of the virulent double-stranded DNA phage PAK_P3 during infection of its host *Pseudomonas aeruginosa* and reveal how its genome rapidly decondenses to adopt a specific 3D organization that reflects its transcriptional program. Concomitantly, the host genome decondenses as gene expression wanes and transcription induced domains vanish. We also uncover specific and discrete bridging of the host and the phage genomes, showing how the phage genome exploits the spatial genome architecture of its host to succeed in its infection cycle. Our data highlights an unprecedented level of genome folding and gene expression regulation during a viral infection.

## Introduction

Mobile genetic elements (MGEs) are ubiquitous and play a major role in shaping the structure, evolution, and function of genomes across all domains of life (Frost et al. 2005; Salmond et Fineran 2015). These elements - including transposons, plasmids, integrons, retroelements, and viruses - are often called genomic “parasites” as they can replicate and move within and between host genomes. Their mobility impacts genome organization (Lawson et al. 2023) and facilitates horizontal gene transfer driving rapid evolution (Khedkar et al. 2022). While genetic and evolutionary consequences of MGEs have been well studied, how MGEs organize themselves into three dimensions and how they influence the physical and spatial organization of host genomes remains understudied.

The advent of proximity ligation technologies (HiC) has transformed our understanding of genome biology (Dekker et al. 2002; Lieberman-Aiden et al. 2009). These techniques allow us to capture how genomes fold into three dimensions inside cells, revealing how chromosomes are organized and how spatial proximity relates to essential processes such as transcription, replication or DNA repair. In eukaryotes, these studies have uncovered principles such as chromatin looping and compartmentalization (Rao et al. 2014; Lieberman-Aiden et al. 2009; Meneu et al. 2025; Conte et al. 2022). In bacteria, HiC has revealed robust structural features like self-interacting domains (CIDs) (Umbarger et al. 2011), alignment of chromosome arms (Wang et al. 2015; Marbouty et al. 2015), links between gene expression and chromatin contacts (Lioy et al. 2018; Bignaud et al. 2024), or *trans* interactions between chromosomes to synchronize their replication (Val et al. 2016). Nevertheless, few studies have investigated the three-dimensional relationships between MGEs and host chromosomes using model organisms (Girard et al. 2025; Moreau et al. 2018; Lamy-Besnier et al. 2023; Kumar et al. 2022; Park et al. 2024). For instance, DNA of the hepatitis B virus associates closely with CpG islands located upstream of active genes in infected human cells, presumably favoring their own replication and gene expression (Moreau et al. 2018). In contrast, yeast *Saccharomyces cerevisiae* 2µ plasmid appears to tether preferentially, but loosely, to regions with low transcriptional activity, potentially to facilitate its hitchhiking segregation strategy (Girard et al. 2025). These few examples suggest that MGEs exploit or adapt to the spatial architecture of host genomes to orchestrate their life cycle.

Among MGEs, virulent phages stand out as they inject their genome into their host to hijack its resources and produce new viral particles that are released upon cell lysis, a process that can be achieved in less than 10 minutes (Dufour et al. 2016). While the transcriptional and biochemical processes taking place during phage infection have been described for many diverse phages (Wicke et al. 2021; Labarde et al. 2021; Blasdel et al. 2017), little is known regarding the structural transitions of phage genomes themselves, from inside the capsid to genomic material delivery into the host, to during infection cycle. Today, the question of whether the genome of virulent phages exploit the spatial architecture of their host to favor their infection strategy, or have selected an independent mechanism, remains open. Similarly, how phage infection influences the spatial genome organization of its host is still unknown.

To explore these questions, we combined time-series HiC with transcriptomics to investigate the genome folding dynamics of the virulent phage PAK_P3 during infection of the opportunistic pathogen *Pseudomonas aeruginosa*. This relatively small (∼88 kb) phage belongs to the *Nankokuvirus* family and was shown to hijack the host RNA polymerase for its own transcription (Chevallereau et al. 2016; Blasdel et al. 2017). With an eclipse period of nearly 12 minutes and an infection cycle duration of around 20 min, phage PAK_P3 is among one of the fastest known dsDNA phages with contractile tails (Chevallereau et al. 2016). By tracking in parallel the variation of phage and bacterial genomes architecture as well as their interactions during infection kinetics, we revealed: (*i*) an overall decondensation of the *P. aeruginosa* chromosome, (*ii*) a rapid and specific dynamics of PAK_P3 genome folding linked to its transcriptional program and, (*iii*) a specific interaction pattern between both genomes, pointing to the existence of dedicated mechanisms set up by the phage to better exploit its host’s cellular machinery and/or the host’s protective response.

By leveraging spatial genomics applied to phage biology, this work sheds light on the mechanisms of phage genome expression, their interactions with hosts, and their subversion mechanisms. The overall results deepen our understanding of phage-host.

## Results

### High resolution chromosome architecture of P. aeruginosa strain PAK

To establish a robust baseline for assessing phage infection-induced chromosomal changes, we first characterized the 3D organization of the genome of *P. aeruginosa* strain PAK (host of the phage PAK_P3) from an exponential culture using a HiC protocol optimized for bacteria (**Methods**). We identified ubiquitous folding features previously reported for model bacteria such as *E. coli*, *B. subtilis* as well as *P. aeruginosa* strain PAO1 (Lioy et al. 2018; 2020; Marbouty et al. 2015; Le et al. 2013): (*i*) a prominent diagonal of strong local contacts, indicative of physical proximity between neighboring loci (**Fig. 1a, Supplementary Fig. 1**); (*ii*) large self-interacting regions dubbed chromosome interaction domains (CIDs, n = 37, **Methods**) whose boundaries correspond to highly transcribed genes (**Fig. 1a**); and (*iii*) a general correlation between gene expression, as assessed from published RNA-seq experiments (Chevallereau et al. 2016), and short-range contacts (Pearson correlation score = +0.69, **Fig. 1a, Methods**) (Lioy et al. 2018; Bignaud et al. 2024). High resolution (500 bp) contact maps highlighted the fine-scale bacterial gene structure, with transcription units enriched in short-range HiC contacts and forming transcription induced domains (TIDs) that interact preferentially with each other (**Fig. 1b**, blue squares), as recently described in *Vibrio cholerae* and *E. coli* (Bignaud et al. 2024). Pileup analysis of averaged contact maps centered on the start codons of the 10% most transcribed transcription units further revealed the same bundle signature at the transcription start sites as previously reported (Bignaud et al. 2024) (**Fig. 1c, Methods**). Interestingly, we also detected a plaid-like pattern at larger scales and visible across the entire contact map, especially when we plotted the autocorrelation score of part of the matrix (**Fig. 1a-d, Methods)**. This pattern, reminiscent of the alternation between transcriptionally active and inactive domains common in eukaryotic genomes (Lieberman-Aiden et al. 2009), will need further analysis to be deciphered and understood in bacteria. Finally, *P. aeruginosa* contact map exhibited a characteristic diagonal perpendicular to the main one, extending from the *parS* cluster near the origin of replication, towards the terminus, reflecting long-range contacts between chromosome replichores, a pattern mediated by SMC-ScpAB complexes (**Fig. 1a, Supplementary Fig. 1**) (Marbouty et al. 2015; Wang et al. 2015; 2017; Lioy et al. 2020). Average 3D projection of the contact map, generated using a probability-based approach based on negative binomial random variables (Varoquaux et al. 2014; 2023; Poinsignon et al. 2023) (**Methods**), recapitulated the global architecture of *P. aeruginosa* chromosome and the cohesion between the two chromosomal replichores (**Fig. 1e**)

**Figure 1:**
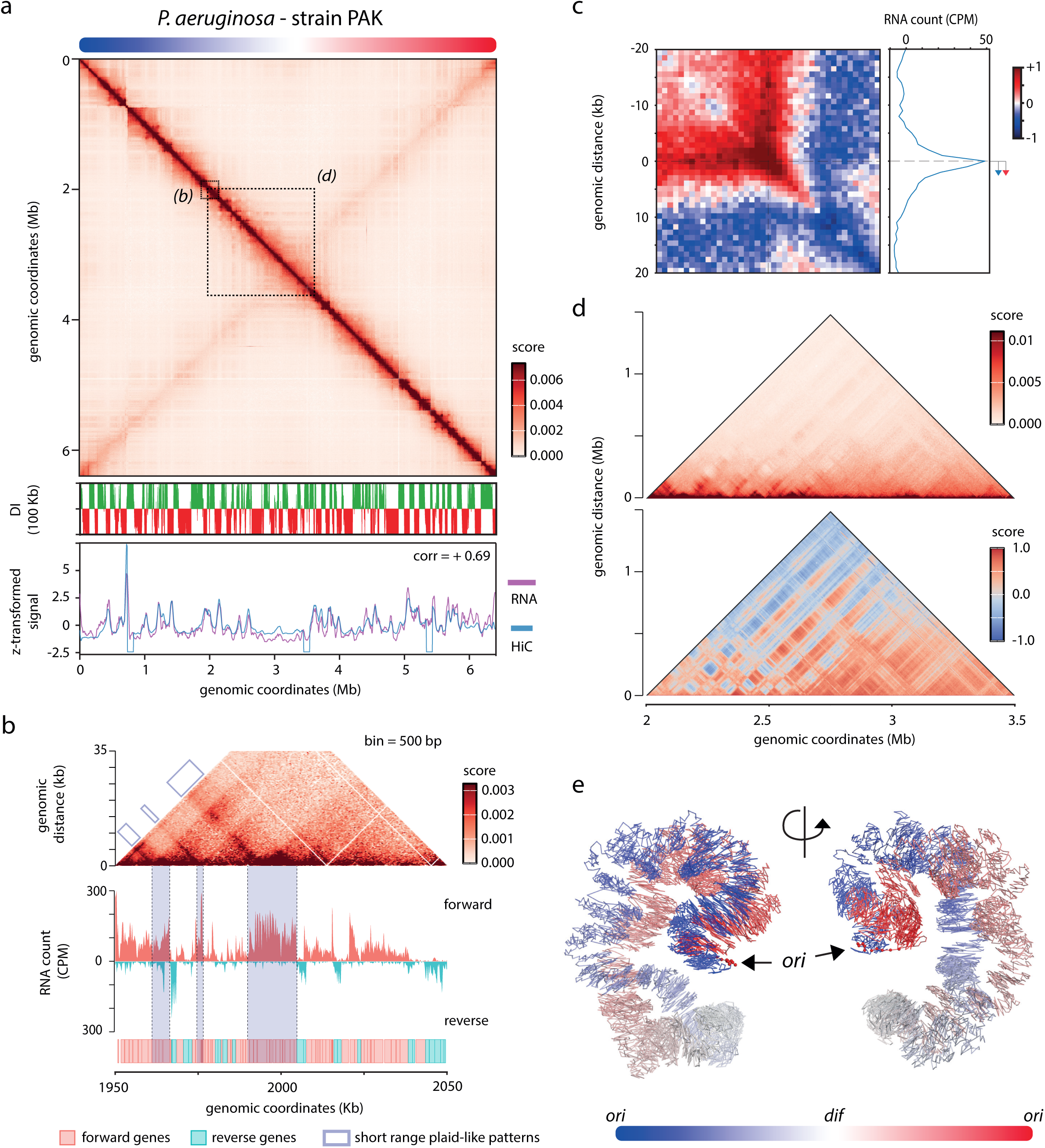
*P. aeruginosa* strain PAK chromosome folding. (a) From top to bottom: 5 kb-resolution HiC contact maps of the strain PAK; directional index at 100 kb; correlation between the DNA short-range contacts (blue – 5kb bins, 25 kb distance) and the transcriptional signal (purple - bin = 5 kb) (Pearson correlation score = +0.78). A linear scheme of the chromosome with a colored gradient (blue – white – red) is indicated above the contact map and refers to the average 3D structure in (e). Dashed squares indicate zoom area for panel (b) and (d) respectively. (b) Magnification of a genomic locus at 500 bp resolution (1.95 – 2.05 Mb). Below the contact map is plotted the RNA signal in count per million (CPM) for the forward (red) and reverse (blue) (bin = 100 bp). The scheme below the plot indicates the orientation of the genes (red = forward, blue = reverse). Blue square indicates examples of compartmented highly transcribed areas (*i.e*. TIDs and plaid-like patterns). (c) Pileup log10 ratio of 20 kb contact map windows (bin = 1 kb) centered on the start codons of the 10% most transcribed transcription units compared to random windows taken along the main diagonal. On the right is the corresponding pileup of transcription tracks (red arrow = forward genes, blue arrow = reverse genes). (d) Zoom of a larger part of the contact map (2 – 3.5 Mb) using either balanced HiC (top) or auto-correlation scores (bottom) (bin = 5kb). (e) Average 3D model of the PAK chromosome. Colors along the structure correspond to the colors of the linear scheme of the chromosome. Origin of replication (*ori*) corresponds to larger red beads and is indicated.

These observations confirmed that HiC captures global and local features of chromosome architecture and was, therefore, suitable to detect potential minimal structural transitions during phage infection.

### PAK_P3 infection leads to a local destructuration of the host chromosome

We performed, in two replicates, HiC on a PAK_P3-infected exponential culture of the bacterial strain PAK at a multiplicity of infection (MOI) of 25 (**Methods**). Given the fast PAK_P3 infection cycle (∼ 20 minutes), we sampled the culture at 0, 3, 5, 7, 10, 13, and 16 minutes post addition of phage (T0 to T16) and processed the cells for HiC (**Methods**). Computation of relative copy number of each chromosome (phage and bacteria) during the time course showed a rapid increase in the number of phage DNA molecules from 0 to 3 min, a relative stabilization between 3 and 7 minutes, followed by a new increase between 7 and 13 minutes (**Supplementary Fig. 2a**). These results were in line with the expected infection cycle of the PAK_P3 phage, with an entry of the phage within the cell population (0-3 min), hijacking of the host (3-7 min) prior to the replication of the phage genome and its encapsidation (7-16 min) (Chevallereau et al. 2016). A comparison of HiC maps recovered from cells along the kinetics showed a progressive decondensation of the host chromosome, accompanied by a disappearance of TIDs and *cis* contacts (**Fig. 2a-b and Supplementary Fig. 2b**) (Bignaud et al. 2024). This phenomenon was clearly observed at 13 min post-infection, where the entire main diagonal appears impacted compared to the non-infected culture (**Fig. 2a-b**). We also observed a loss of local borders, as well as of other 3D structures such as stripes and loops (**Fig. 2b)**. In contrast, the secondary diagonal signal was retained, suggesting a persistence of replichores bridging (**Fig. 2a and Supplementary Fig. 2-3**) and that the SMC-ScpAB complex is not impacted by the infection at this stage. Similarly, the average 3D structure of the PAK genome during the infection reflected the structural changes inferred from the HiC map: a conservation of the global organization accompanied by a disruption of local folding of the chromosome (**Fig. 2c**).

**Figure 2:**
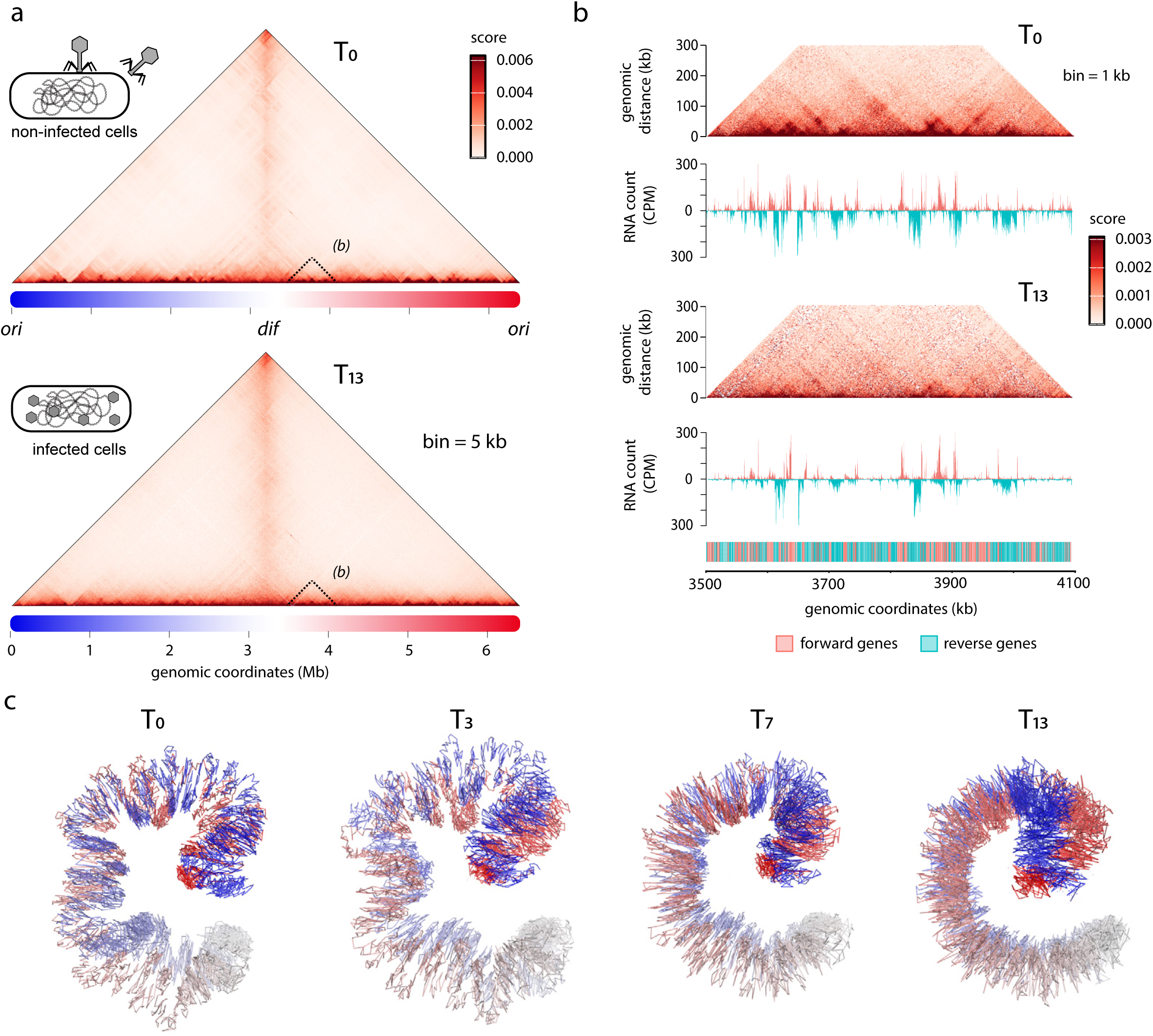
Impact of phage PAK_P3 infection on *P. aeruginosa* chromosome architecture. (a). Contact maps at 5 kb resolution at T0 (top) and T13 (bottom). A linear scheme of the chromosome with a colored gradient (blue – white – red) is indicated below the contact maps and refers to the average 3D structure in (c). (b) Magnification of a genomic locus (3.5 Mb - 4.1 Mb) at T0 and T13 revealing the impact of phages on genome architecture at short scale (bin = 1 kb). Below the two contact maps are plotted the RNA count in CPM for the forward (red) and reverse (blue) (bin = 100 bp). Map of the genes (blue = forward, red = reverse). Genomic coordinates of the studied loci are indicated below. The different scale bars are indicated on the right. (e) Average 3D models of the PAK chromosome at 0, 3, 7 and 13 minutes after addition of phage PAK_P3. The color of the structure is the same as in Fig.1e.

These observations, reminiscent of the effect of rifampicin on bacterial chromosome architecture (Bignaud et al. 2024; Marbouty et al. 2015), agreed with a progressive arrest of the host transcription concomitant to the hijacking of the transcriptional machinery by phage PAK_P3 (Chevallereau et al. 2016).

### PAK_P3 genome rapidly decondenses and reorganizes following its transcriptional program

In addition to providing information about the host’s genome organization, HiC also revealed the dynamic changes experienced by the PAK_P3 genome during the infection time course. We then analyzed the architecture of the phage genome for each timepoint, from T0, when the phage is still inside its capsid, until T16 at the end of the infection (**Fig. 3 and Supplementary Fig. 4**). At T0, the contact decay curve or genomic distance law p(s) was nearly flat, suggesting relatively equal contacts between pairs of DNA segments independently of the distance separating them (**Fig. 3a** – orange curve). This is consistent with a highly condensed and constrained genome, where DNA is tightly packed and with no preferred structure as shown by the contact map and the average 3D projection of the phage genome (**Fig. 3b**). In contrast, the p(s) curve transitions from flat to sharply declining during the first time points, demonstrated a rapid decondensation of the viral chromosome as it enters the host cytoplasm, with a minimum reached between 5 and 10 min (**Fig. 3a**). Then, the p(s) gradually switched back towards a flat state, and at 16 min it had nearly returned to its initial T0 state, suggesting that most of the newly synthesized PAK_P3 genomes were encapsidated into formed viral particles. Comparison between transcriptional profiles (Chevallereau et al. 2016) and contact maps generated over the course of infection further highlighted the dynamic structural changes experienced by the phage genome (**Fig. 3c-d**). Segments of the genome indeed showed an increase in medium range contacts that correlated with regions carrying genes expressed at early, middle and late stages of infection (**Fig. 3c-d**). As these domains do not correlate with specific GC content, a potential bias of HiC experiments, it’s likely that the structures observed with HiC were linked to transcription of the phage genome (**Supplementary Fig. 5**). In addition, at 3 minutes and in both replicates, a 14 kb region encompassing part of the early expressed genes adopted a hairpin-like structure and isolated itself from the rest of the genome (**Fig. 3d-e and Supplementary Figure 5**). This discrete structure was also conveniently illustrated by the average 3D reconstruction of the phage genome at 3 minutes after infection (**Fig. 3f** – dashed circle) and points to the possible action of one or more specific factors involved in structuring phage DNA. The tip of the hairpin consisted in two sequences devoid of genes and transcription (**Fig. 3e –** dashed circles point to the tip of the structure and dashed areas to the anchors sequences). This phenomenon suggests that phages genomes are actively structured like cellular organisms. We also observed a variation of the overall circular signal all along the phage cycle with a clear increase between 5 and 10 minutes followed by a decrease at 13 minutes (**Fig. 3a-d**). This circular-like structure as well as the decondensation - recondensation of the phage genome was also observed on the 3D representation of the HiC data generated all along the phage infection cycle (**Fig. 3d**). These observations suggested two potential types of replication of the phage DNA already described for Myoviridae (« Myoviridae | ICTV », s. d.): a rolling circle replication that generates concatenates of phage genomes or a theta replication that solely produces circular molecules. Analysis of the HiC coverage showed a clear skew from both extremities to the center between 7 and 10 minutes supporting the existence of circular molecules and the theta replication hypothesis associated with a single start site for replication (**Fig. 3d**).

**Figure 3:**
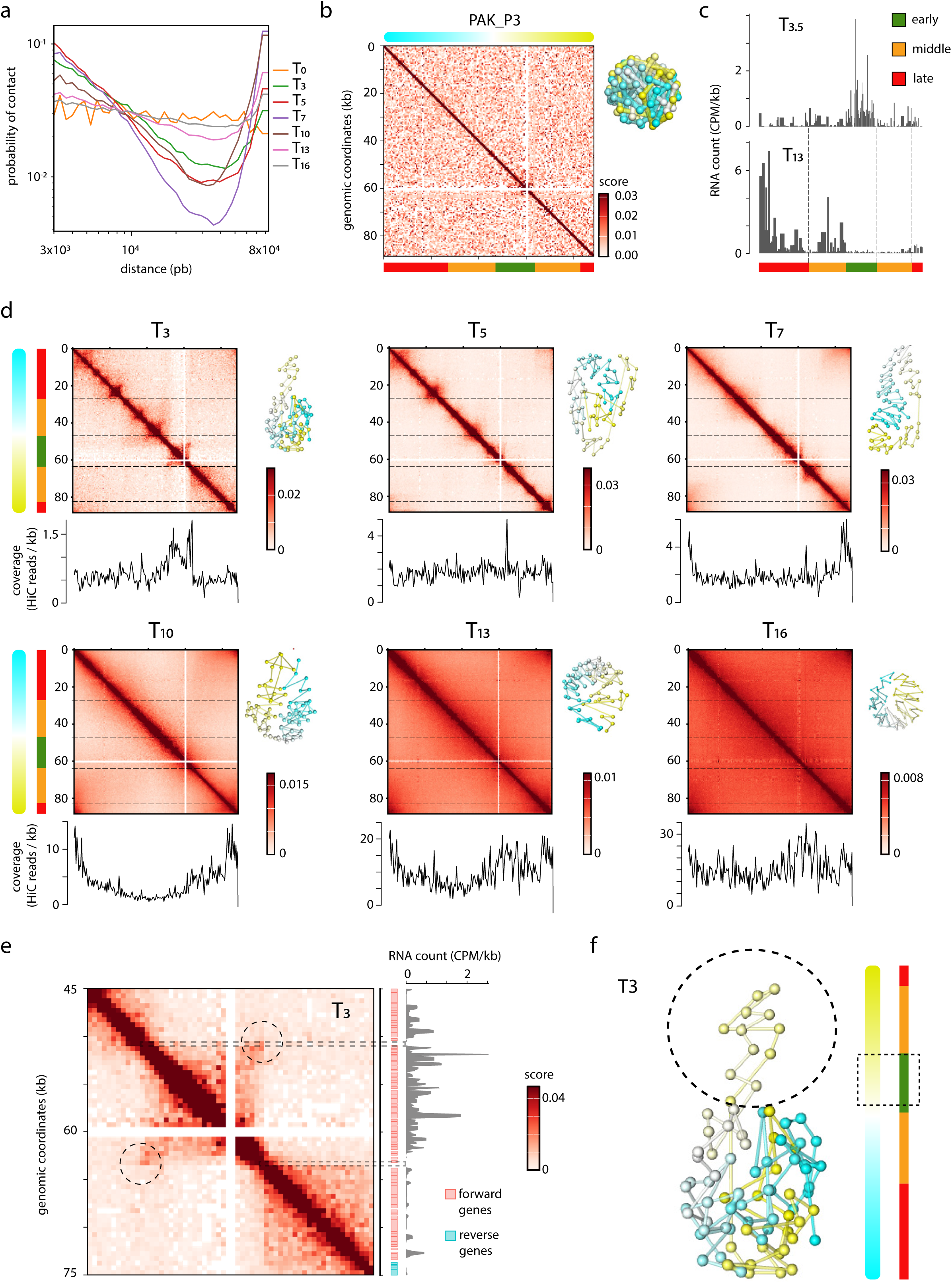
Dynamics of phage PAK_P3 genome during infection. (a) Normalized contact in function of the genomic distance for the PAK_P3 genome at the different time points (p(s)). Normalized scores and genomic coordinates are in log scale. The color of each time point is indicated on the right. (b) Contact map of PAK_P3 (bin = 500 bp) after adding of the phage particles to the cell culture (T0). A linear colored scheme (cyan – white – yellow) of the PAK_P3 genome is indicated above the contact map and refers to the phage average 3D projection of the HiC data (on the right of the map). A linear colored scheme (red, orange, green) is drawn below the map and refers to the transcriptional program of the phage (c). (c) RNA count for T3.5 and T13 from previous publication (Chevallereau et al. 2016). Colored segments below the plot indicate the different gene clusters corresponding to early (green), middle (orange) and late (red) transcribed genes). Dashed lines correspond to the borders of the early, mid and late clusters respectively. (d) Contact maps of PAK_P3 for time points 3, 5, 7, 10, 13 and 16 minutes. A 3D projection of the HiC data for each time point is plotted on the right of the corresponding contact maps. The HiC coverage is also plotted below each map (bin = 1kb). (e) magnification of the contact map at the locus encompassing the hairpin-like structure (45kb-75kb). RNA count (T = 3.5 min) is plotted on the right of the map. Forward (red) and reverse (blue) genes are also indicated. Grey area points to the two anchors loci positioned at the tip of the structure. (f) 3D model of phage PAK_P3 genome at 3 min post addition of the phage. The color of the structure is the same as for (a). The dashed circle and square indicate the loop/hairpin seen on the HiC map.

Overall, our data showed that the temporal regulation of PAK_P3 genome structure is strongly associated with the carefully regulated activation of its gene expression program, starting at the earliest stages of infection (3 min).

### PAK_P3 exhibits a specific interaction pattern with the host chromosome

We next computed the contacts between the phage DNA with the host chromosome at each time point during phage infection (**Fig. 4a and Supplementary Fig. 6**) (**Methods**). The normalized score of contacts between PAK_P3 DNA molecules and the host chromosome increased from 0 to 7 min before decreasing until 16 min, in agreement with the packaging of the phage between 7 and 16 minutes isolating its DNA from the host. We also detected specific, enriched contacts of the phage with the host’s chromosome (**Fig. 4a-b**, examples of valley and peak of interactions are labelled as areas I and II). These contacts suggested a specific underlying mechanism that may favor hijacking of the host transcriptional machinery. To test whether such contacts corresponded to specific localization of the phage genome and/or a more generic increased accessibility of the host nucleoprotein filament by episomes, we performed a HiC experiment on the *P. aeruginosa* strain PAK carrying the multicopy 6kb plasmid pJN105. The 4C-like contacts plot between plasmid and the bacterial chromosome displayed a pattern inversely correlated to the phage contacts plot (**Fig. 4b-c**, Pearson correlation score: −0.356), demonstrating different mechanisms of interaction of these two genomic entities with the host DNA and, therefore, different localizations within cellular space. We then asked if these different patterns could be linked to the HiC signal at short range or to host transcription activity, two signals known to be correlated (**Fig. 1a**). A plot of the contacts between PAK_P3 and *P. aeruginosa* chromosome after 5 minutes of infection regarding HiC signal at the same time or RNA profile at 3.5 minutes of infection hinted that PAK_P3 interact preferentially with weakly transcribed regions of the chromosome (Pearson correlation score = −0.4257 and −0.2573 respectively, **Fig. 4b-c**). On the contrary, the pJN105 4C-like contacts plot exhibits a clear correlation with these two signals (Pearson correlation score = +0.6723 and +0.4123 respectively, **Fig. 4b-c**). To go further in this analysis, we computed the Pearson correlation score between contact score and HiC signal as well as transcriptomic data for all time points and showed that, in contrast to plasmid pJN105, phage genomes contacts during the first half of the infection cycle show an anti-correlation with HiC signal and transcriptional profile of the host chromosome (**Fig. 4d and Supplementary Fig. 6**).

**Figure 4:**
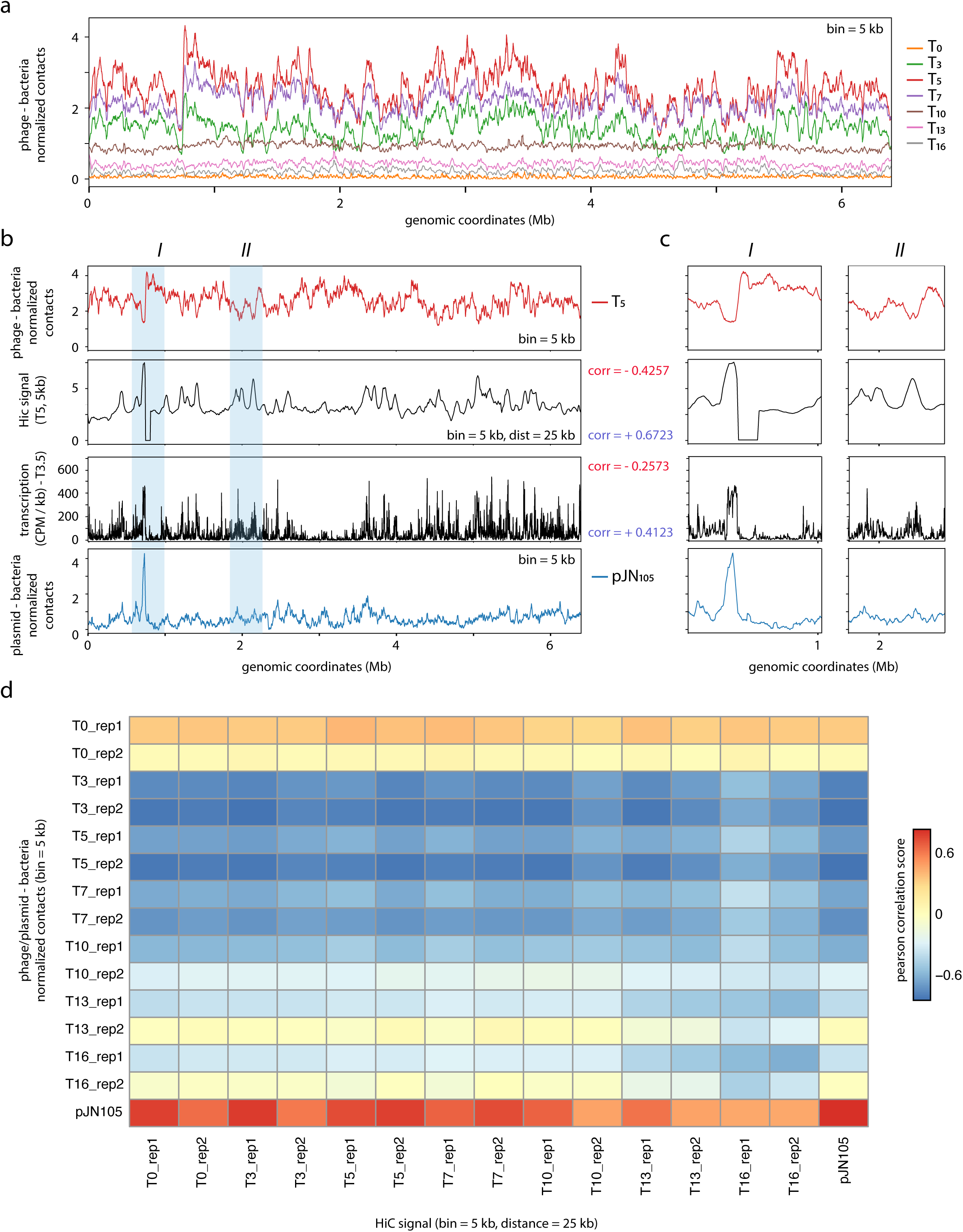
interactions between PAK_P3 and *P. aeruginosa* chromosome. (a) 4C-like contact of PAK_P3 genome with the PAK genome (5 kb bins). Colors are the same as for Figure 3 and are indicated on the right of the graph. (b, c) From top to bottom: 4C-like contact (5 kb bins) of PAK_P3 genome with PAK genome after 5 minutes of infection, HiC signal at short range (distance = 25 kb, bin = 5 kb), transcription signal of the PAK genome at 3$‘.5 minutes in CPM per kilobase, 4C-like contact (5 kb bins) of plasmid pJN105 with the PAK genome. Genomic coordinates of the PAK genome are indicated at the bottom of the plots. Pearson correlation coefficients between 4C-like contact profiles (red = PAK_P3 at T5; blue = pJN105) and either HiC or RNA signal is given on the right of the panel. Two magnifications of the profiles are plotted in (c) (area I and II indicated in blue in (b)). (d) Matrix indicating the Pearson correlation score between the 4C-like contact of PAK_P3 or pJN105 genome with the PAK genome and the HiC signal at short range for the same time points (distance = 25 kb, bin = 5 kb) at the different time points.

Overall, the HiC temporal analysis of the infection cycle of phage PAK_P3 revealed an unexpected behavior of its genome that interacts preferentially with low active host chromosome loci, unlike a high-copy plasmid. Whether such a profile is due to an active positioning of the phage genome or is an indirect consequence of either the phage or the host metabolism remains to be elucidated.

## Discussion

Since the discovery of phages more than a century ago, research on these highly diverse viruses has yielded fundamental insights into a broad range of processes related to DNA metabolism (Salmond et Fineran 2015). Several studies have begun to decipher the mechanism of action of bacterial infection by phages, notably in *P. aeruginosa* (Chevallereau et al. 2016; Blasdel et al. 2017; Wicke et al. 2021; Chaikeeratisak et al. 2017). In the present work, we have deciphered the infection cycle of the virulent *Pseudomonas* phage PAK_P3 using proximity-ligation-based methods. This pioneering work shed light on new processes concerning its genome organization, its dynamics, as well as its impact on the architecture of the *P. aeruginosa* chromosome and offer valuable insights into the complex interplay between this phage and its bacterial host (**Fig. 5**).

**Figure 5:**
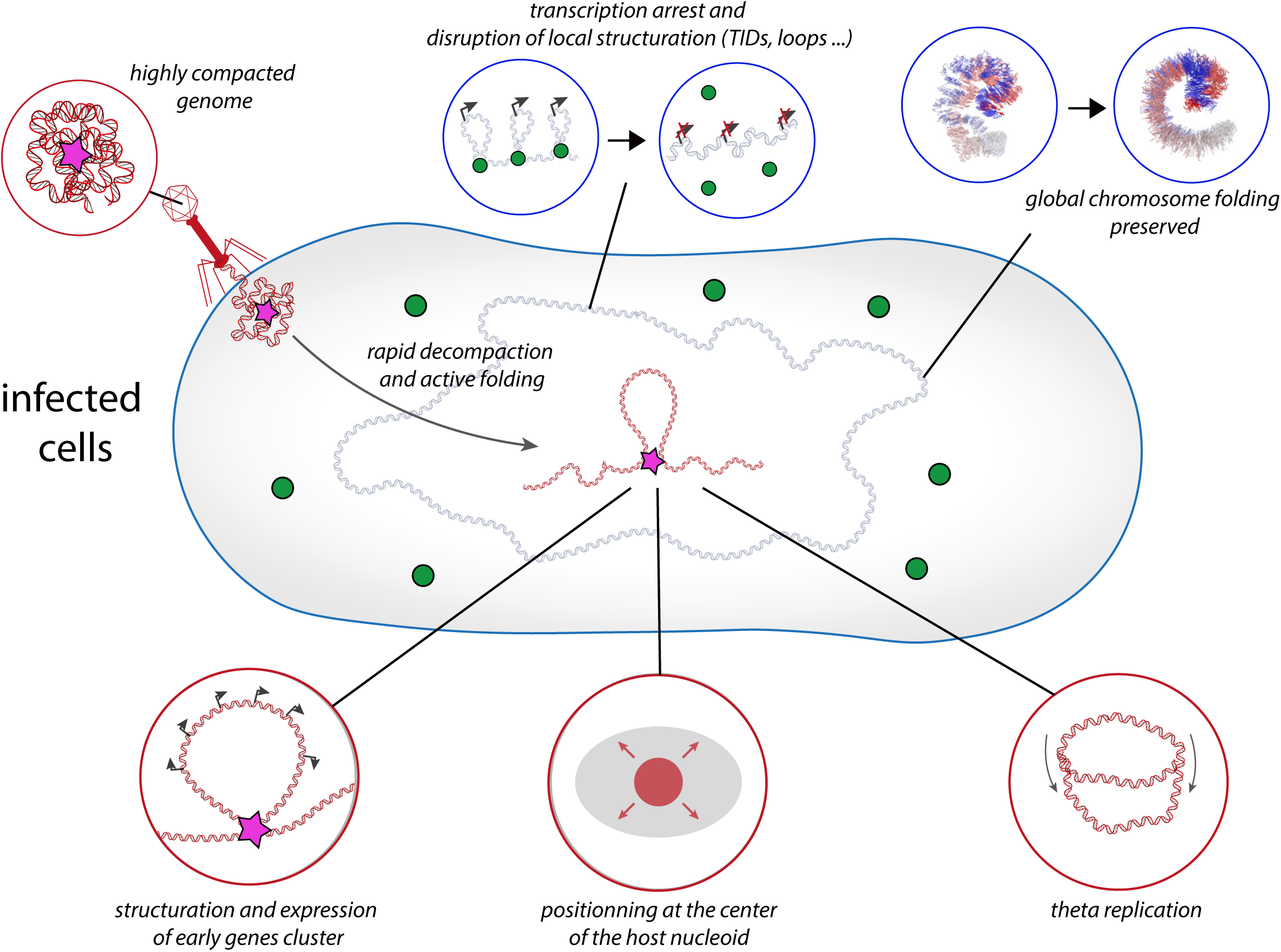
model of PAK_P3 genome dynamics and impact on host chromosome during its infection cycle in *P. aeruginosa*. Red and blue DNA molecules correspond to phage and host genomes respectively. Green circles refer to host proteins while pink star refers to NAP-like factor present in the viral particles and responsible for the early structuration of the phage genome. Black arrows correspond to transcription.

First, PAK_P3 infection led to a rapid decondensation of the host chromosome, loss of local bundle correlations with transcription, and alterations in chromatin structures. These observations are reminiscent of the effects of rifampicin on bacterial chromosome architecture (Marbouty et al. 2015; Lioy et al. 2018) and confirm a gradual arrest of host transcription due to phage hijacking of the transcriptional machinery as previously demonstrated using transcriptomic data (Chevallereau et al. 2016).

The temporal analysis of phage infection reported also provides new critical insights into the dynamics of phage genomes during infection. The rapid decondensation of its genome upon entry into the host cytoplasm, followed by a recompaction phase demonstrates dynamic changes in its structure during infection. This dynamic genome restructuring notably led to the formation of structural domains whose borders correlate with the phage transcriptional program. One striking observation comes from the early time point (T3) where the locus encompassing early expressed genes adopts a hairpin-like structure (**Fig 3 and Fig. 5**). This result points to an active process that raises the question of the existence of NAP-like (Nucleoid Associated Proteins) factors or a non-coding RNA that could actively and rapidly structure the phage genome. This could imply that such a factor may be encapsidated with the phage genome (**Fig. 5**, pink star). Interestingly, among the peptides identified from the proteomic analysis of PAK_P3 virions (Henry et al. 2013), few matched Gp160 (YP_008857795.1), a protein that is predicted to belong to the family of short DNA binding protein, as all known bacterial NAPs (**Supplementary Table 2, Methods**). Moreover, this factor is encoded by a mono-cistronic gene and its expression is not temporally regulated during the infection (Chevallereau et al. 2016). To strengthen the hypothesis of the involvement of specific phage proteins in the early structuration of the phage genome, additional studies using phage mutants and/or Chip-seq experiments could be engaged. Nevertheless, we already demonstrate that phage genomes, as the genomes of cellular organisms, can be highly structured during their reproductive cycle.

Our study also reveals that PAK_P3 phage interacts preferentially with specific locations of the *P. aeruginosa* genome. The fact that PAK_P3 and the multicopy plasmid pJN105 exhibit opposite interaction patterns demonstrates that MGEs have different ways of taking advantage of the host genome architecture. In the case of PAK_P3, our data shows that it interacts with low transcriptionally active areas. Interestingly, some studies on bacteria have shown that active chromatin tends to localize at the periphery of the nucleoid (Yang et al. 2019; Bignaud et al. 2024). Therefore, our results suggest that PAK_P3 locates its genome at the center of the nucleoid, a location that may be favorable for hijacking the transcription machinery of *P. aeruginosa*. The development of dedicated tools for imaging PAK_P3 genome in infected cells would be required to provide additional evidence supporting the above hypothesis.

In summary, our study highlights the infection strategy employed by a virulent phage infecting *P. aeruginosa* and uncovers intricate interactions between viral and host genomes. These findings open new avenues for further exploration into the mechanisms underlying phage infection and their impact on host chromosome architecture. Understanding these processes at a molecular level will provide an advanced comprehension of virus-host interactions and may open new avenues for developing phage-based therapies.

## Methods

### Bacteriophage and Bacterial strains

Strains (PAK and PAK + pJN105) were grown in Lennox broth (LB) or on Lennox agar at 37°C. PAK_P3 phage was amplified in exponential growing cultures of strain PAK for approximately 6 hours. Bacterial debris was removed first by centrifugation at 5200 g for 10 min at 4°C then cell lysate supernatants were 0.22 µm filtered and kept at 4°C. Phage stock concentrations were then quantified by spot dilution assay (Henry et al. 2013).

### Kinetics of phage infection

Strain PAK was grown until mid-exponential phase (OD = 0.3) in 80 mL of LB. PAK_P3 phage was then added at a MOI of 25 (Chevallereau et al. 2016). 8 mL aliquots of the phage bacterial culture were removed at specific intervals of the phage bacterial infection cycle: 0, 3, 5, 7, 10, 13, 16 minutes. 1 mL was used to perform a spot assay and quantify phage loads at the different intervals of infection. The remaining 7 mL was immediately transferred into 15 mL falcon tubes containing 1 mL of Formaldehyde 35.5-37% to perform HiC experiments. Tubes were then incubated for 30 min under shaking at RT, followed by incubation at 4°C for another 30 min. Formaldehyde was quenched by adding 2 mL of Glycine 2.5 M followed by an incubation of 20 min under shaking. Pellets were recovered by a centrifugation of 10 min at 10,000 x g at 4°C, washed in PBS and then re-centrifuged using the same settings. Pellets were then frozen in dry ice and stored at −80°C until use.

### HiC libraries

Proximity-ligation libraries were generated using the ARIMA kit (Arima Genome-Wide HiC Kit). Samples were first incubated for 1 h on ice and then resuspended in 1 mL of sterile water and transferred to 2 mL precellys tubes containing glass beads of 0.1- and 0.5-mm diameter (Precellys – Bertin Technology). Cells were disrupted using the precellys apparatus (Precellys Evolution) and the following program (7500 rpm, 6 cycles 30 sec ON / 30 sec OFF, 4°C). Tubes were then centrifuged for 1 min at 1000 g and 700 µL of lysate was recovered and transferred to a new 1.5 mL eppendorf tube. Tubes were centrifuged for 20 min at 16,000 g, 4°C. Supernatant was carefully removed and pellet was resuspended in 45 µL of water. We then followed the ARIMA protocol with the addition of a centrifugation step after the digestion (yellow solution) as previously described (Bignaud et al. 2025). Briefly, after the digestion step, tubes were centrifuged for 20 min at 16,000 g, 4°C. Supernatant was carefully removed and pellet was resuspended in 100 µL of water before applying the next step of the kit (blue solution) and following the protocol to the end. DNA was then purified using AMPure beads and eluted in 130 µL of water before processing for sequencing.

### Libraries processing and sequencing

HiC genomic libraries were first sheared at a mean size of 500 bp using a covaris S200 apparatus and then processed on streptavidin beads using the colibri kit (ThermoFisher) as previously described (Cockram et al. 2021). Libraries were sequenced on Nextseq or NovaSeq apparatus.

### HiC data analysis

Contact maps were generated using hicstuff (Matthey-Doret et al. 2020) and pre-digested reads (bowtie2 - very sensitive local mode – mapping quality of 30) and a reference FastA files containing the phage PAK_P3 (or the plasmid pJN105) and the PAK genome. Contact maps were then binned at different resolutions (500 bp, 1 kb, 2 kb and 5 kb), balanced using cooler (Abdennur et Mirny 2019), analyzed and displayed using the R package “HiCexperiments”) (Serizay et al. 2024). All the scripts for processing the data are available at the following link: https://github.com/mmarbout/PAK_P3_infection_analysis.

### RNAseq analysis

RNA raw fastq files (Chevallereau et al. 2016) were recovered from SRA database and processed using homemade script (tinymapper - https://github.com/js2264/tinyMapper). Processed RNA files for phage transcription signal were also recovered from the corresponding study (NCBI GEO portal - GSE76513). RNA data were combined with HiC using different R packages and plotted using R.

### 3D models

Mcool files were used as input for the workflow 3D genome builder (https://github.com/data-fun/3d-genome-builder). PDB files obtained were then modified using custom script and visualized using pymol.

### Prediction of nucleoid-associated proteins (NAPs)

Using HMMsearch (hmmer.org), we searched for PAK_P3 proteins homologous to 14 known bacterial nucleoid-associated proteins (NAPs) and histones (HU, H-NS, Lsr2, HMF, Hc1, Fis, IHF-A, IHF-B, Dps, Hfp, MvaT, CbpA, XrvA and the bacterial histones), but detected no significant matches. We next screened PAK_P3 proteins with HMMsearch against PFAM-A (release 35.0) and selected domains annotated in pfam2go (version 2025/04/29, (Mitchell et al. 2015)) as ‘DNA-binding’ or ‘transcription factor’ domains. Only YP_008857685.1 (gp50), containing a predicted DNA_pol_A domain, was identified. Finally, based on nucleoid-associated protein (NAP) criteria (Hocher et al. 2022), proteins from phage PAK_P3 were screened for short length (≤300 amino acids), predicted DNA-binding activity, and monocistronic gene organization. Size-filtered proteins were first analysed using DNA Binder (Kumar et al. 2007), followed by Operon Mapper (Taboada et al. 2018). Ten proteins were retained and subsequently examined for their presence in PAK_P3 virions (Henry et al. 2013) (**Supplementary Table 2**).

### Statistics & Reproducibility

No data were excluded from the analyses. Statistical tests were done using R.

## Data availability

Sequencing data have been deposited in the NCBI under the BioProject number PRJNA1331554. Mcool, bigwig, bed and pdb files are available on Zenodo (10.5281/zenodo.17529627). All of the sequencing data and their corresponding access number used in the present study are listed in Supplementary Table 1.

## Code availability

All the codes used for the analysis are available at the following address https://github.com/mmarbout/PAK_P3_infection_analysis.

## Acknowledgements

We thank Jacques Serizay for his help in analyzing the data and his critical comments on our manuscript.

This research was supported by funding to MM and LD from PRCI ANR-20-CE92-0048 (PhaStGut project); to RK from the European Research Council under the Horizon 2020 Program (ERC grant agreement 771813). AB is supported by an ENS fellowship by the French Ministry of Higher Education, Research and Innovation. DEC, QLB and FG were funded through the PhastGut ANR grant.

## Authors’ contributions

Conceptualization: MM, AB, QLB, LD. Data curation: AB, MM. Formal analysis: PM, AB, MM. Funding acquisition: RK, LD, MM. Investigation: AB, MM, LD, RK. Methodology: QLB, AT, DEC, FG, MM. Project administration: LD, RK, MM. Resources: LD, MM. Software: PM, AB, MM. Supervision: LD, MM. Validation: RK, LD, MM. Visualization: RK, LD, MM. Writing - editing: all authors.

## Rights and permissions

A CC-BY public copyright license has been applied by the authors to the present document and will be applied on all subsequent versions up to the Author Accepted Manuscript arising from this submission, in accordance with the grants’ open access conditions.

## Supplementary data

**Supplementary Figure 1:**
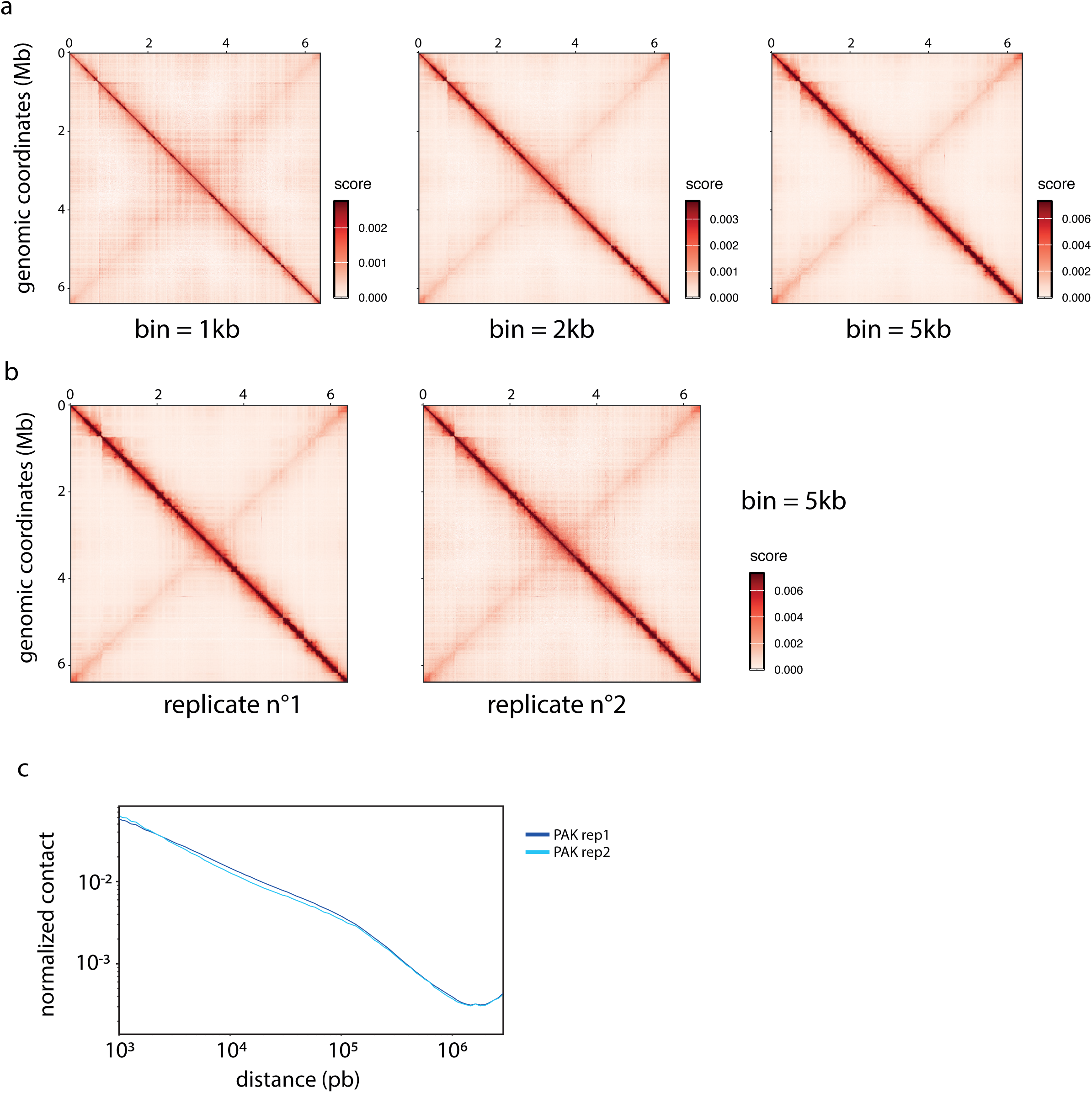
contact map at various resolutions and of the replicates. (a) From left to right: 1 kb-resolution, 2 kb-resolution and 5 kb-resolution Hi-C contact maps of the strain PAK. Genomic coordinates are indicated on the left and on the top of the contact maps while scale bars on the right. (b) 5 kb-resolution Hi-C contact maps of the two replicates for the strain PAK. (c) P(s) contact law for the two replicates.

**Supplementary Figure 2:**
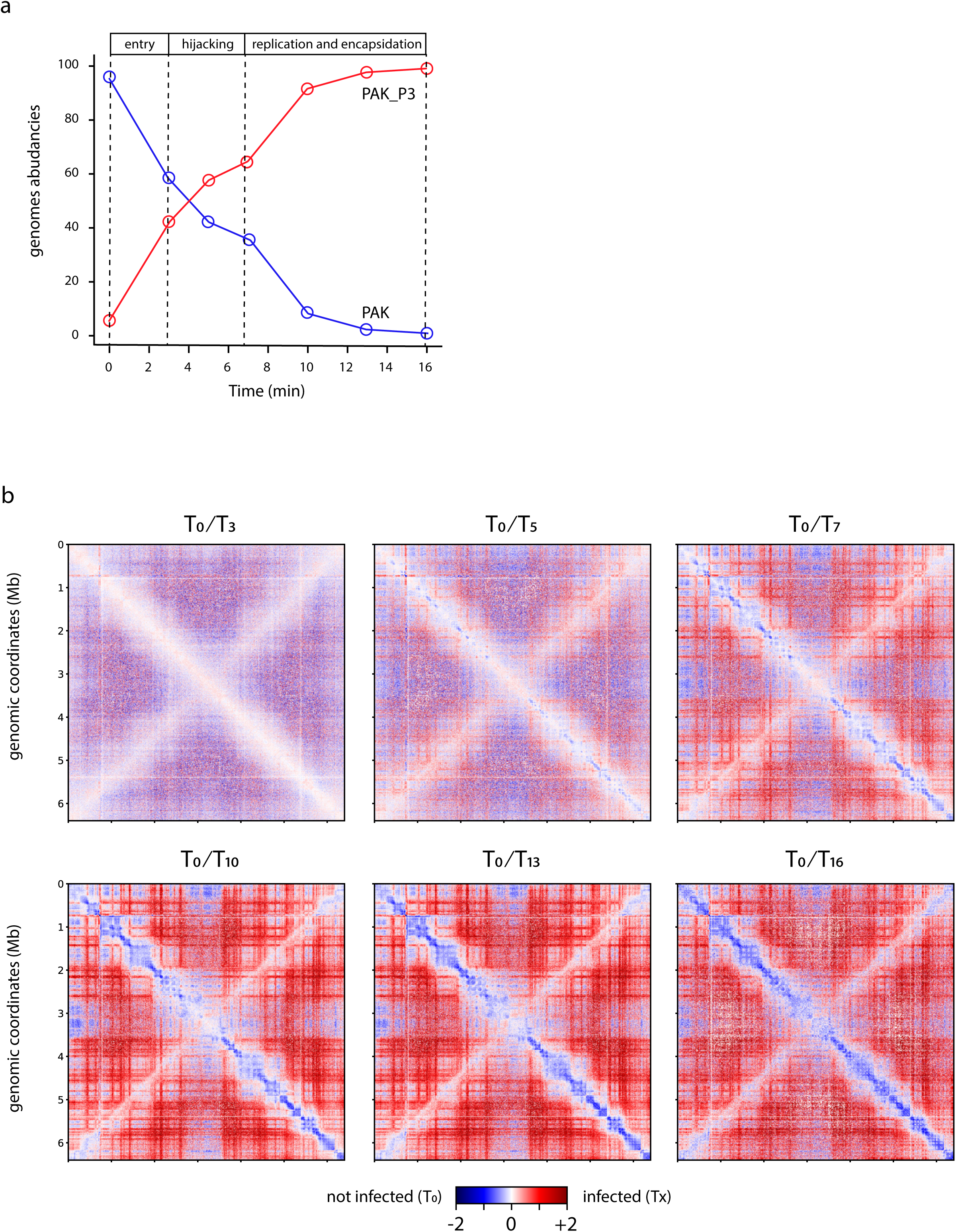
replicates of PAK_P3 infection cycle. (a) Graphic of PAK (blue) and PAK_P3 (red) genomes abundances in function of time. (b) 5 kb-resolution ratio contact maps of the strain PAK during the PAK_P3 time course infection cycle. Each map is a comparison between a defined time point and the time 0 minute. Genomic coordinates are indicated on the map. A log scale bar is indicated on the right of the contact maps.

**Supplementary Figure 3:**
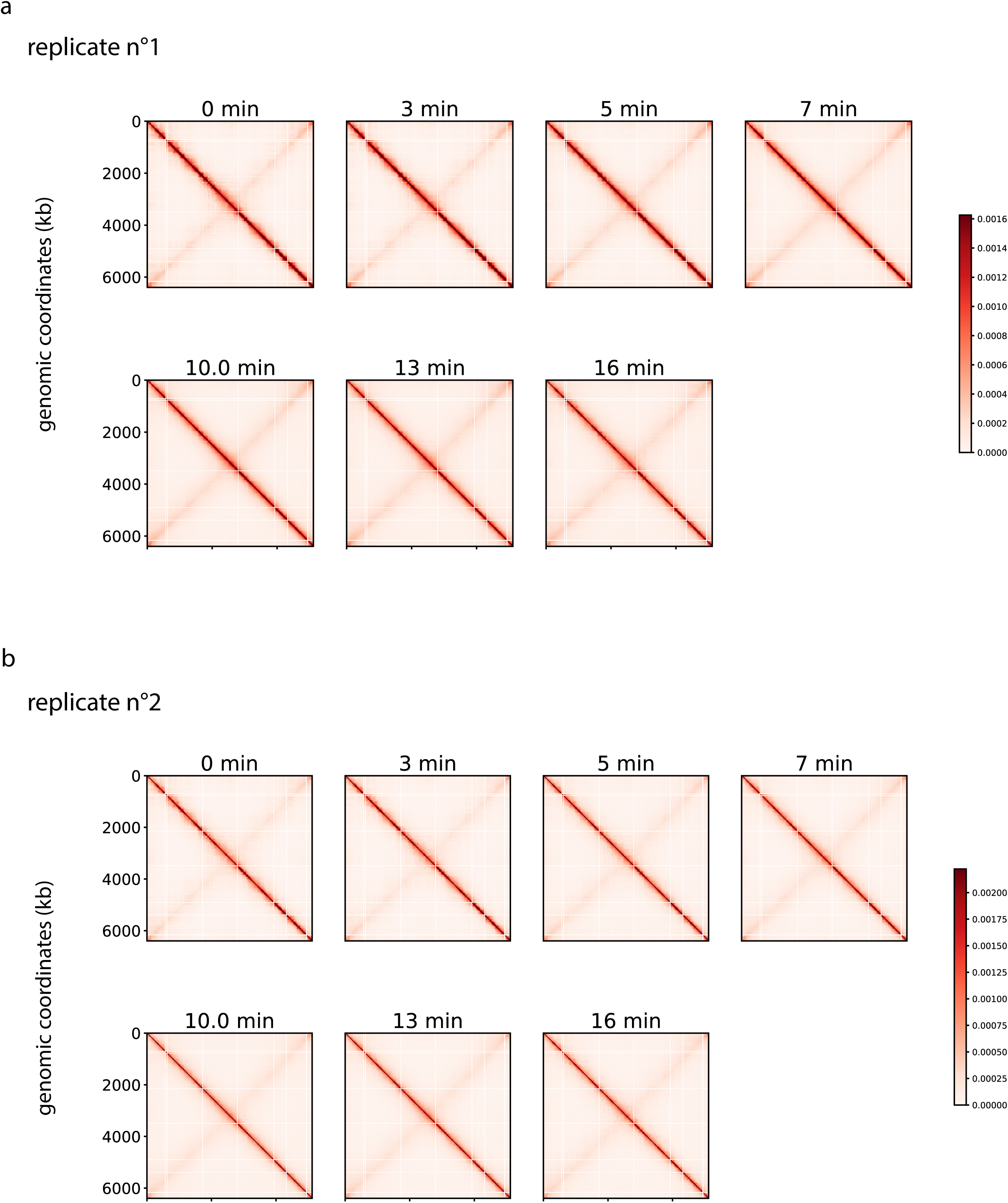
contact map of replicates of PAK during infection cycle. 5 kb-resolution contact maps of strain PAK for time points 0, 3, 5, 7, 10, 13 and 16 minutes (bin = 5kb). Genomic coordinates are indicated on the left of the maps. (a) Replicate 1. (b) Replicate 2.

**Supplementary Figure 4:**
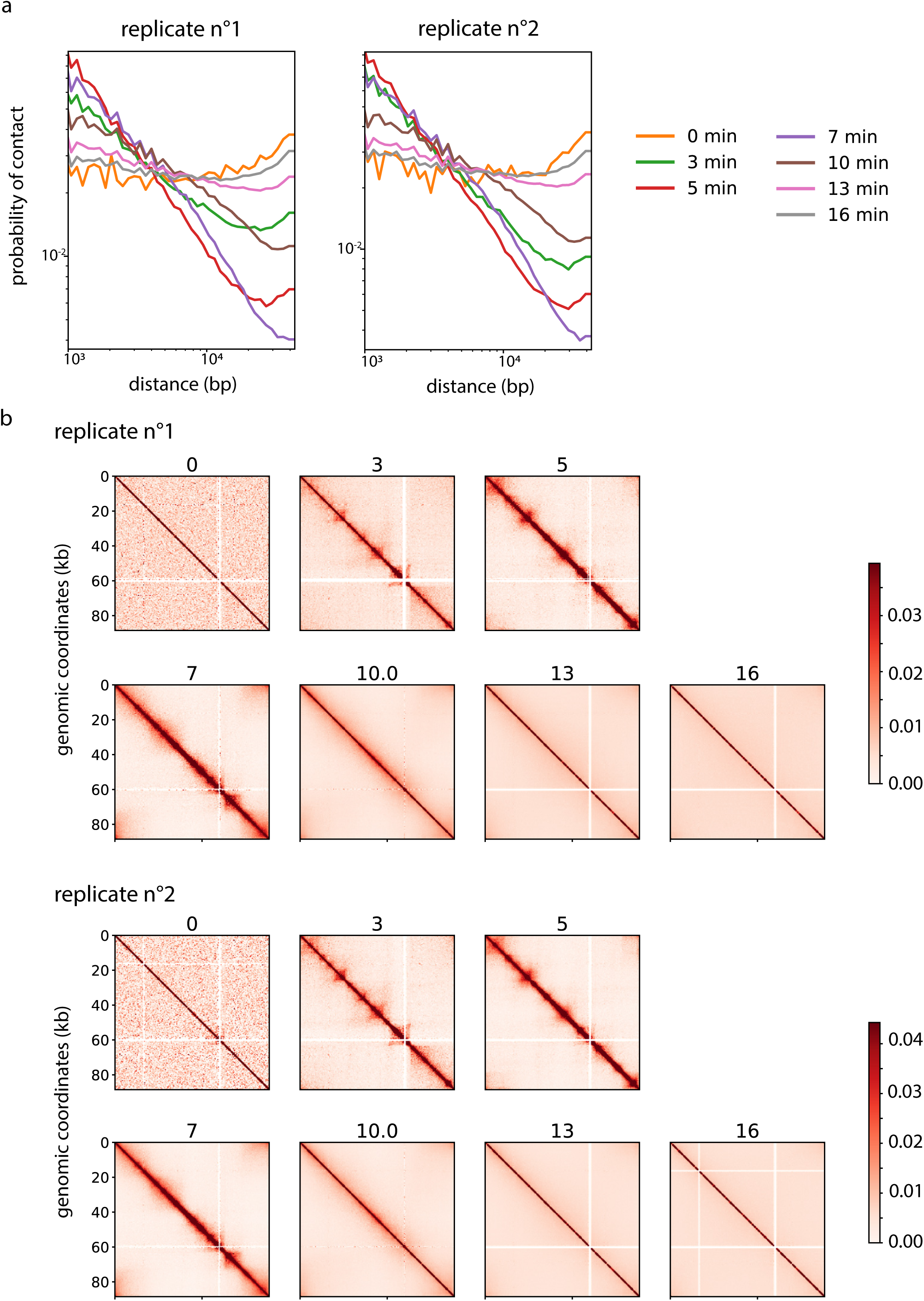
contact map of replicates of PAK_P3 during infection cycle. (a) Normalized contact in function of the genomic distance for the PAK_P3 genome at the different time points and for the two replicates. Normalized scores and genomic coordinates are in log scale. The color of each time point is indicated on the right. (b) Contact maps of PAK_P3 for time points 0, 3, 5, 7, 10, 13 and 16 minutes (bin = 500 pb). Genomic coordinates are indicated on the left of the contact map. Top: replicate 1; bottom: replicate 2. Panels on the right corner indicate the GC content of the PAK_P3 genome (bin = 1kb).

**Supplementary Figure 5:**
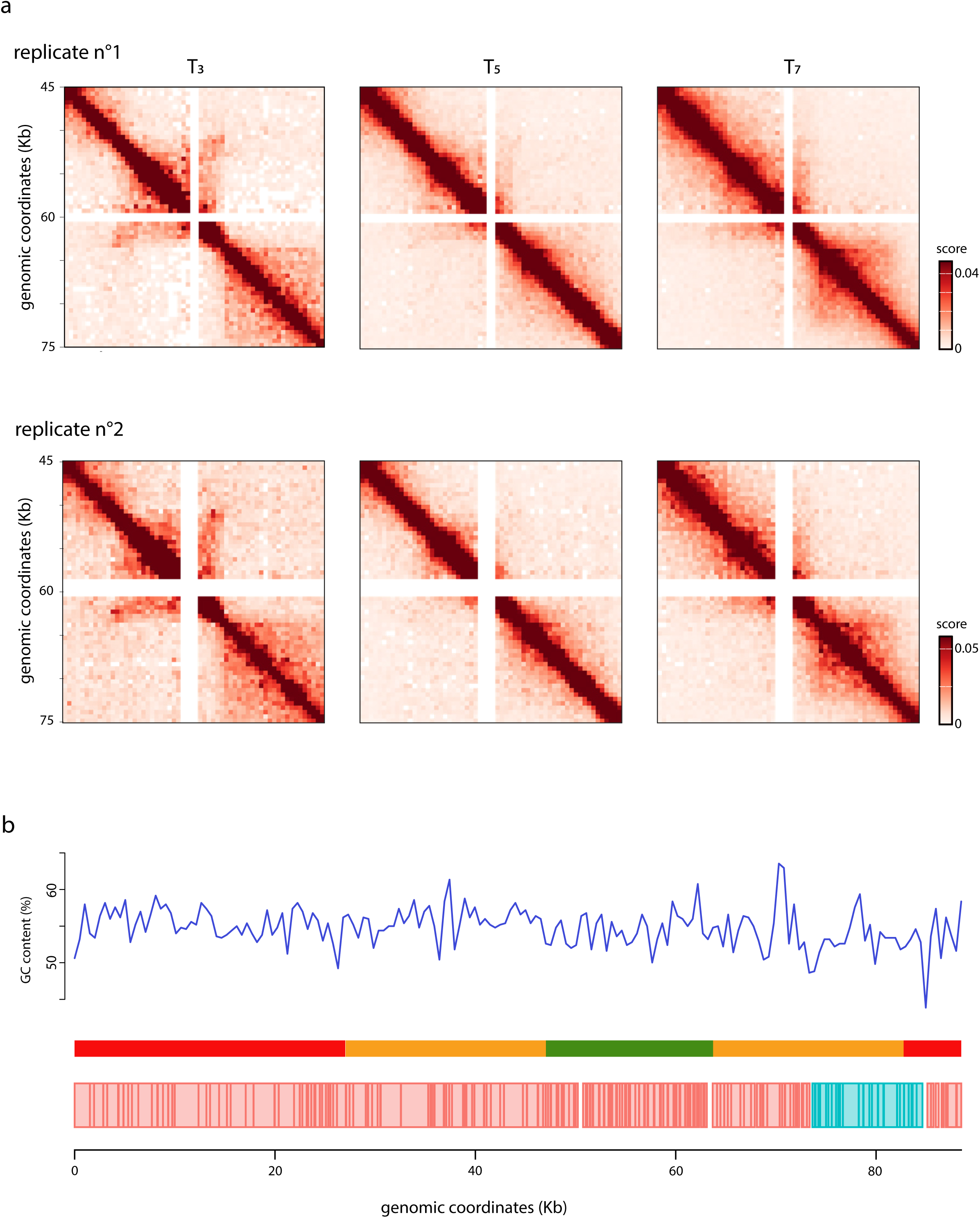
contact maps of PAK_P3 hairpin locus and GC content of the PAK_P3 genome. (a) 500 pb-resolution of magnification of Hi-C contact maps of the PAK_P3 at 3, 5 and 7 minutes between 45 to 75 kb. Genomic coordinates are indicated on the map. Scale bar is indicated on the right. Top: replicate 1; bottom: replicate 2. (b) GC content along the PAK_P3 genome (bin = 1kb). Schemes of the cluster (early, middle, late) and the genes (forward and reverse are indicated below the plot.

**Supplementary Figure 6:**
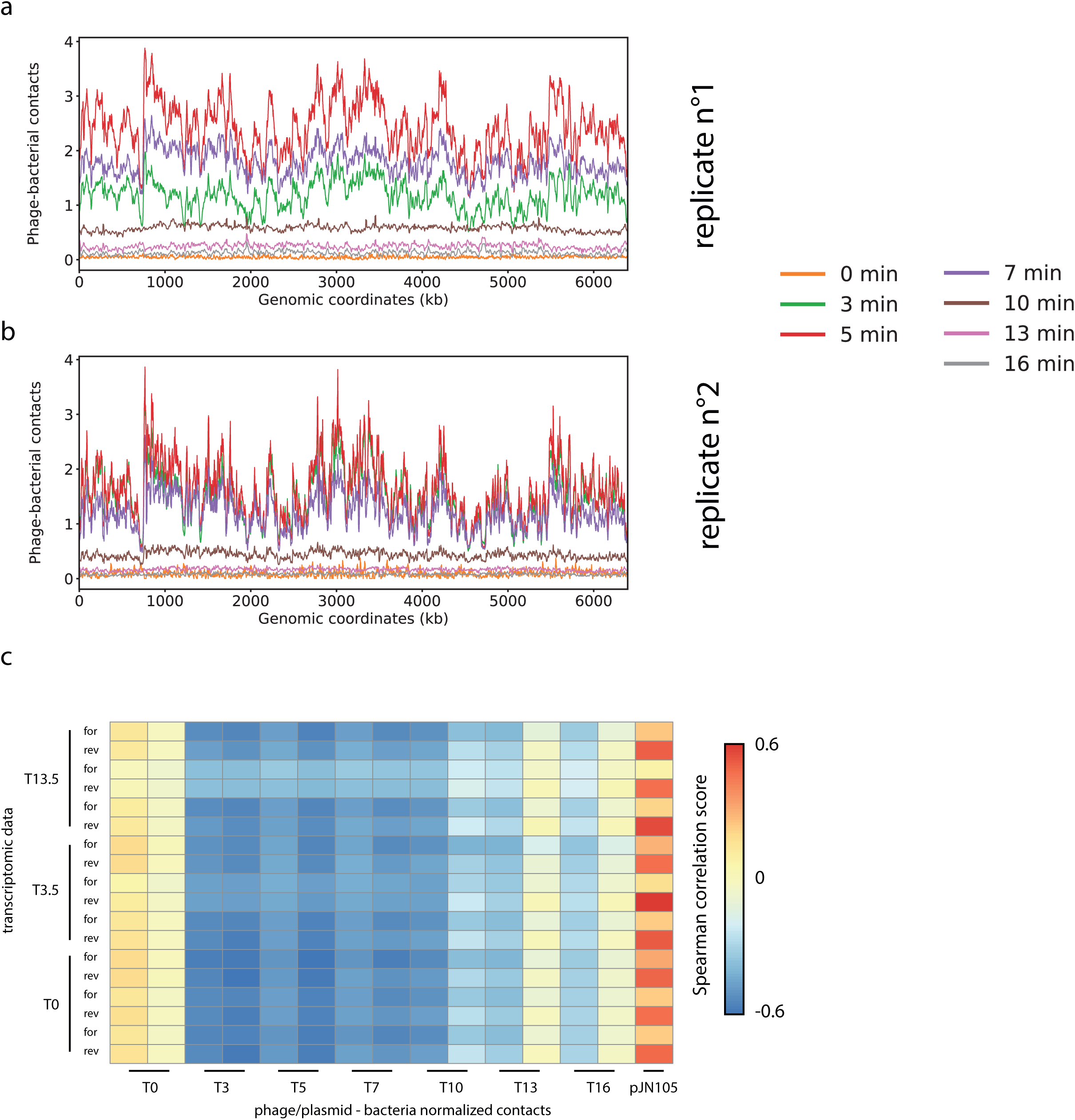
contact of PAK_P3 with PAK genome. (a, b) 4C-like contact (10 kb bins) of PAK_P3 genome with the PAK genome at the different time points and for the two replicates (a - replicate 1; b - replicate 2). Genomic coordinates of the PAK genome are indicated below the graph. (c) matrix indicating the Pearson correlation score between the 4C-like contact of PAK_P3 genome or pJN105 with the PAK genome at the different time points (1 kb bin) and the transcriptomic signal at T0, T3.5 and T13.

**Supplementary Table 1:**
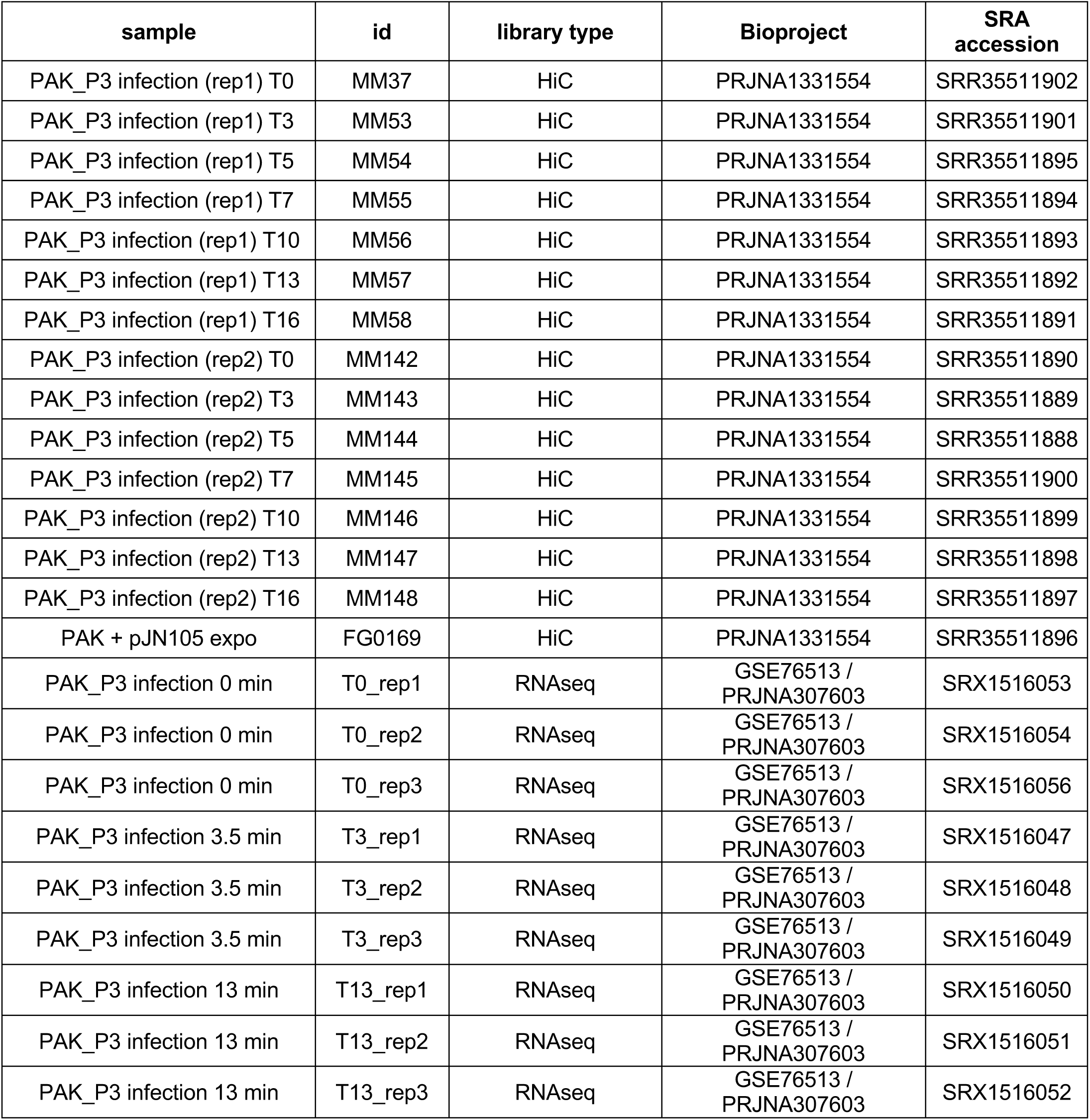
datasets used in the present study.

**Supplementary Table 2:** NAPs candidate proteins encoded in PAK_P3 genome. (* Henry *et al*., 2013). The predicted function corresponds to the function written in the GFF file.

| Protein ID | Protein name | Length (aa) | Predicted function (GFF) | presence in PAK_P3 particles * |
| --- | --- | --- | --- | --- |
| YP_008857687.1 | gp52 | 61 | Protein of unknown function | No |
| YP_008857715.1 | gp80 | 70 | Protein of unknown function | No |
| YP_008857729.1 | gp94 | 78 | Protein of unknown function | No |
| YP_008857733.1 | gp98 | 81 | Protein of unknown function | No |
| YP_008857735.1 | gp100 | 171 | Protein of unknown function | No |
| YP_008857741.1 | gp106 | 152 | Protein of unknown function | No |
| YP_008857772.1 | gp137 | 76 | Protein of unknown function | No |
| YP_008857795.1 | gp160 | 58 | Protein of unknown function | Yes |
| YP_008857798.1 | gp163 | 100 | Protein of unknown function | No |
| YP_008857799.1 | gp164 | 272 | Terminase large subunit | No |

## Bibliography

Abdennur, Nezar, et Leonid A Mirny. 2019. « Cooler: scalable storage for Hi-C data and other genomically labeled arrays ». Bioinformatics 36 (1): 311–16. 10.1093/bioinformatics/btz540.

Bignaud, Amaury, Charlotte Cockram, Céline Borde, et al. 2024. « Transcription-Induced Domains Form the Elementary Constraining Building Blocks of Bacterial Chromosomes ». Nature Structural & Molecular Biology, janvier 4, 1–9. 10.1038/s41594-023-01178-2.

Bignaud, Amaury, Devon E. Conti, Agnès Thierry, et al. 2025. « Phages with a Broad Host Range Are Common across Ecosystems ». Nature Microbiology, septembre 19, 1–13. 10.1038/s41564-025-02108-2.

Blasdel, Bob G., Anne Chevallereau, Marc Monot, Rob Lavigne, et Laurent Debarbieux. 2017. « Comparative Transcriptomics Analyses Reveal the Conservation of an Ancestral Infectious Strategy in Two Bacteriophage Genera ». The ISME Journal 11 (9): 1988–96. 10.1038/ismej.2017.63.

Chaikeeratisak, Vorrapon, Katrina Nguyen, MacKennon E. Egan, Marcella L. Erb, Anastasia Vavilina, et Joe Pogliano. 2017. « The Phage Nucleus and Tubulin Spindle Are Conserved among Large Pseudomonas Phages ». Cell Reports 20 (7): 1563–71. 10.1016/j.celrep.2017.07.064.

Chevallereau, Anne, Bob G. Blasdel, Jeroen De Smet, et al. 2016. « Next-Generation “-Omics” Approaches Reveal a Massive Alteration of Host RNA Metabolism during Bacteriophage Infection of Pseudomonas Aeruginosa ». PLOS Genetics 12 (7): e1006134. 10.1371/journal.pgen.1006134.

Cockram, Charlotte, Agnès Thierry, et Romain Koszul. 2021. « Generation of Gene-Level Resolution Chromosome Contact Maps in Bacteria and Archaea ». STAR Protocols 2 (2): 100512. 10.1016/j.xpro.2021.100512.

Conte, Mattia, Ehsan Irani, Andrea M. Chiariello, et al. 2022. « Loop-Extrusion and Polymer Phase-Separation Can Co-Exist at the Single-Molecule Level to Shape Chromatin Folding ». Nature Communications 13 (1): 4070. 10.1038/s41467-022-31856-6.

Dekker, Job, Karsten Rippe, Martijn Dekker, et Nancy Kleckner. 2002. « Capturing Chromosome Conformation ». Science 295 (5558): 1306–11. 10.1126/science.1067799.

Dufour, Nicolas, Olivier Clermont, Béatrice La Combe, et al. 2016. « Bacteriophage LM33_P1, a Fast-Acting Weapon against the Pandemic ST131-O25b:H4 Escherichia Coli Clonal Complex ». The Journal of Antimicrobial Chemotherapy 71 (11): 3072–80. 10.1093/jac/dkw253.

Frost, Laura S., Raphael Leplae, Anne O. Summers, et Ariane Toussaint. 2005. « Mobile Genetic Elements: The Agents of Open Source Evolution ». Nature Reviews Microbiology 3 (9): 722–32. 10.1038/nrmicro1235.

Girard, Fabien, Antoine Even, Agnès Thierry, et al. 2025. « Parasitic Plasmids Are Anchored to Inactive Regions of Eukaryotic Chromosomes through a Nucleosome Signal ». The EMBO Journal 44 (7): 2134–56. 10.1038/s44318-025-00389-1.

Henry, Marine, Rob Lavigne, et Laurent Debarbieux. 2013. « Predicting In Vivo Efficacy of Therapeutic Bacteriophages Used To Treat Pulmonary Infections ». Antimicrobial Agents and Chemotherapy 57 (12): 5961–68. 10.1128/aac.01596-13.

Hocher, Antoine, Guillaume Borrel, Khaled Fadhlaoui, Jean-François Brugère, Simonetta Gribaldo, et Tobias Warnecke. 2022. « Growth Temperature and Chromatinization in Archaea ». Nature Microbiology 7 (11): 1932–42. 10.1038/s41564-022-01245-2.

Khedkar, Supriya, Georgy Smyshlyaev, Ivica Letunic, et al. 2022. « Landscape of mobile genetic elements and their antibiotic resistance cargo in prokaryotic genomes ». Nucleic Acids Research 50 (6): 3155–68. 10.1093/nar/gkac163.

Kumar, Ashish, Yuanzhi Lyu, Yuichi Yanagihashi, et al. 2022. « KSHV Episome Tethering Sites on Host Chromosomes and Regulation of Latency-Lytic Switch by CHD4 ». Cell Reports 39 (6). 10.1016/j.celrep.2022.110788.

Kumar, Manish, Michael M. Gromiha, et Gajendra P. S. Raghava. 2007. « Identification of DNA-Binding Proteins Using Support Vector Machines and Evolutionary Profiles ». BMC Bioinformatics 8 (novembre): 463. 10.1186/1471-2105-8-463.

Labarde, Audrey, Lina Jakutyte, Cyrille Billaudeau, et al. 2021. « Temporal Compartmentalization of Viral Infection in Bacterial Cells ». Proceedings of the National Academy of Sciences of the United States of America 118 (28): e2018297118. 10.1073/pnas.2018297118.

Lamy-Besnier, Quentin, Amaury Bignaud, Julian R. Garneau, et al. 2023. « Chromosome folding and prophage activation reveal specific genomic architecture for intestinal bacteria ». Microbiome 11 (1): 111. 10.1186/s40168-023-01541-x.

Lawson, Heather A., Yonghao Liang, et Ting Wang. 2023. « Transposable Elements in Mammalian Chromatin Organization ». Nature Reviews Genetics 24 (10): 712–23. 10.1038/s41576-023-00609-6.

Le, Tung B K, Maxim V Imakaev, Leonid A Mirny, et Michael T Laub. 2013. « High-Resolution Mapping of the Spatial Organization of a Bacterial Chromosome ». Science (New York, N.Y.) 342 (6159): 731–34. 10.1126/science.1242059.

Lieberman-Aiden, Erez, Nynke L. van Berkum, Louise Williams, et al. 2009. « Comprehensive Mapping of Long-Range Interactions Reveals Folding Principles of the Human Genome ». Science 326 (5950): 289–93. 10.1126/science.1181369.

Lioy, Virginia S., Axel Cournac, Martial Marbouty, et al. 2018. « Multiscale Structuring of the E. Coli Chromosome by Nucleoid-Associated and Condensin Proteins ». Cell 172 (4): 771–783.e18. 10.1016/j.cell.2017.12.027.

Lioy, Virginia S., Ivan Junier, Valentine Lagage, Isabelle Vallet, et Frédéric Boccard. 2020. « Distinct Activities of Bacterial Condensins for Chromosome Management in Pseudomonas Aeruginosa ». Cell Reports 33 (5): 108344. 10.1016/j.celrep.2020.108344.

Marbouty, Martial, Antoine Le Gall, Diego I. Cattoni, et al. 2015. « Condensin- and Replication-Mediated Bacterial Chromosome Folding and Origin Condensation Revealed by Hi-C and Super-Resolution Imaging ». Molecular Cell 59 (4): 588–602. 10.1016/j.molcel.2015.07.020.

Matthey-Doret, Cyril, baudrly, Amaury, et al. 2020. koszullab/hicstuff: Use miniconda layer for docker and improved P(s) normalisation. Zenodo, released octobre 5. 10.5281/zenodo.4066363.

Meneu, Léa, Christophe Chapard, Jacques Serizay, et al. 2025. « Sequence-dependent activity and compartmentalization of foreign DNA in a eukaryotic nucleus ». Science 387 (6734): eadm9466. 10.1126/science.adm9466.

Mitchell, Alex, Hsin-Yu Chang, Louise Daugherty, et al. 2015. « The InterPro Protein Families Database: The Classification Resource after 15 Years ». Nucleic Acids Research 43 (Database issue): D213–221. 10.1093/nar/gku1243.

Moreau, Pierrick, Axel Cournac, Gianna Aurora Palumbo, et al. 2018. « Tridimensional Infiltration of DNA Viruses into the Host Genome Shows Preferential Contact with Active Chromatin ». Nature Communications 9 (1): 4268. 10.1038/s41467-018-06739-4.

« Myoviridae | ICTV ». s. d. Consulté le 28 novembre 2025. https://ictv.global/report_9th/dsDNA/Myoviridae.

Park, Kiwon, Dohoon Lee, Jiseok Jeong, Sungwon Lee, Sun Kim, et Kwangseog Ahn. 2024. « Human Immunodeficiency Virus-1 Induces Host Genomic R-Loop and Preferentially Integrates Its Genome near the R-Loop Regions ». eLife 13 (novembre). 10.7554/eLife.97348.2.

Poinsignon, Thibault, Mélina Gallopin, Pierre Grognet, Fabienne Malagnac, Gaëlle Lelandais, et Pierre Poulain. 2023. « 3D models of fungal chromosomes to enhance visual integration of omics data ». NAR Genomics and Bioinformatics 5 (4): lqad104. 10.1093/nargab/lqad104.

Rao, Suhas S. P., Miriam H. Huntley, Neva C. Durand, et al. 2014. « A 3D Map of the Human Genome at Kilobase Resolution Reveals Principles of Chromatin Looping ». Cell, publication en ligne anticipée, décembre 11. 10.1016/j.cell.2014.11.021.

Salmond, George P. C., et Peter C. Fineran. 2015. « A Century of the Phage: Past, Present and Future ». Nature Reviews Microbiology 13 (12): 777–86. 10.1038/nrmicro3564.

Serizay, Jacques, Cyril Matthey-Doret, Amaury Bignaud, Lyam Baudry, et Romain Koszul. 2024. « Orchestrating Chromosome Conformation Capture Analysis with Bioconductor ». Nature Communications 15 (1): 1072. 10.1038/s41467-024-44761-x.

Taboada, Blanca, Karel Estrada, Ricardo Ciria, et Enrique Merino. 2018. « Operon-Mapper: A Web Server for Precise Operon Identification in Bacterial and Archaeal Genomes ». Bioinformatics (Oxford, England) 34 (23): 4118–20. 10.1093/bioinformatics/bty496.

Umbarger, Mark A, Esteban Toro, Matthew A Wright, et al. 2011. « The Three-Dimensional Architecture of a Bacterial Genome and Its Alteration by Genetic Perturbation ». Molecular Cell 44 (2): 252–64. 10.1016/j.molcel.2011.09.010.

Val, Marie-Eve, Martial Marbouty, Francisco de Lemos Martins, et al. 2016. « A checkpoint control orchestrates the replication of the two chromosomes of Vibrio cholerae ». Science Advances 2 (4). 10.1126/sciadv.1501914.

Varoquaux, Nelle, Ferhat Ay, William Stafford Noble, et Jean-Philippe Vert. 2014. « A statistical approach for inferring the 3D structure of the genome ». Bioinformatics 30 (12): i26–33. 10.1093/bioinformatics/btu268.

Varoquaux, Nelle, William S Noble, et Jean-Philippe Vert. 2023. « Inference of 3D genome architecture by modeling overdispersion of Hi-C data ». Bioinformatics 39 (1): btac838. 10.1093/bioinformatics/btac838.

Wang, Xindan, Hugo B. Brandão, Tung B. K. Le, Michael T. Laub, et David Z. Rudner. 2017. « Bacillus subtilis SMC complexes juxtapose chromosome arms as they travel from origin to terminus ». Science 355 (6324): 524–27. 10.1126/science.aai8982.

Wang, Xindan, Tung B. K. Le, Bryan R. Lajoie, Job Dekker, Michael T. Laub, et David Z. Rudner. 2015. « Condensin Promotes the Juxtaposition of DNA Flanking Its Loading Site in Bacillus Subtilis ». Genes & Development 29 (15): 1661–75. 10.1101/gad.265876.115.

Wicke, Laura, Falk Ponath, Lucas Coppens, Milan Gerovac, Rob Lavigne, et Jörg Vogel. 2021. « Introducing Differential RNA-Seq Mapping to Track the Early Infection Phase for Pseudomonas Phage ɸKZ ». RNA Biology 18 (8): 1099–110. 10.1080/15476286.2020.1827785.

Yang, Sora, Seunghyeon Kim, Dong-Kyun Kim, et al. 2019. « Transcription and Translation Contribute to Gene Locus Relocation to the Nucleoid Periphery in E. Coli ». Nature Communications 10 (1): 5131. 10.1038/s41467-019-13152-y.

